# Gut metabolites influence susceptibility of neonatal mice to cryptosporidiosis

**DOI:** 10.1101/2020.09.11.294462

**Authors:** Kelli L. VanDussen, Lisa J. Funkhouser-Jones, Marianna E. Akey, Deborah A. Schaefer, Kevin Ackman, Michael W. Riggs, Thaddeus S. Stappenbeck, L. David Sibley

## Abstract

The protozoan parasite *Cryptosporidium* is a leading cause of diarrheal disease in those with compromised or under-developed immune systems, particularly infants and toddlers in resource-poor localities. As an enteric pathogen, *Cryptosporidium* invades the apical surface of intestinal epithelial cells, where it resides in close proximity to metabolites in the intestinal lumen. However, the effect of gut metabolites on susceptibility to *Cryptosporidium* infection remains largely unstudied. Here, we first identified which gut metabolites are prevalent in neonatal mice when they are most susceptible to *Cryptosporidium parvum* infection, and then tested the isolated effects of these metabolites on *C. parvum* invasion and growth. Our findings demonstrate that medium or long-chain saturated fatty acids inhibit *C. parvum* growth, while long-chain unsaturated fatty acids enhance *C. parvum* invasion. The influence of these two classes of metabolites on *C. parvum* infection likely reflects the streamlined metabolism in *C. parvum*, which is unable to synthesize fatty acids. Hence, gut metabolites, either from diet or produced by the microbiota, play an important role in the early susceptibility to cryptosporidiosis seen in young animals.

**Importance:** *Cryptosporidium* occupies a unique intracellular niche that exposes the parasite to both host cell contents and the intestinal lumen, including metabolites from the diet and produced by the microbiota. Both dietary and microbial products change over the course of early development, and could contribute to the changes seen in susceptibility to cryptosporidiosis in humans and mice. Consistent with this model, we show that the immature gut metabolome influenced growth of *C. parvum in vitro* and may increase susceptibility to infection in young mice. Interestingly, metabolites that significantly altered parasite growth were fatty acids, a class of molecules that *Cryptosporidium* is unable to synthesize de novo. The enhancing effects of polyunsaturated fatty acids and the inhibitory effects of saturated fatty acids provide further insight into reliance on fatty acid salvage and metabolism of this enteric parasite.

## Main

*Cryptosporidium* has gained notoriety in recent years due to its surprising prevalence as a major enteric diarrheal pathogen in children under two years of age in Africa and South East Asia(1, 2). The parasite is transmitted by a direct oral-fecal route, often through the ingestion of environmentally resistant oocysts in contaminated water supplies(3). Cryptosporidiosis in humans is primarily caused by two species: *Cryptosporidium parvum* infects a wide variety of domestic livestock and is transferred to humans as a zoonotic infection, although some subtypes are known to circulate more directly between humans(4, 5). By contrast, *C. hominis* is almost exclusively transmitted human-to-human(5, 6). Treatment options for cryptosporidiosis are very limited as the only FDA-approved drug, nitazoxanide, is ineffective in immunocompromised patients and not approved for use in children(7).

Numerous studies demonstrate that neonatal animals are highly susceptible to *Cryptosporidium* and that resistance to infection increases with age in mice(8), dairy calves(9), and humans(1, 2). In fact, the Global Enteric Multicenter Study (GEMS) found that, in developing countries, *Cryptosporidium* was the second leading cause of diarrheal episodes in infants (0-11 months of age), the third leading cause in toddlers (12 – 23 months of age) and nearly absent in children two years and older(1, 2). Why neonatal animals are particularly susceptible to the parasite, and what causes them to become resistant as they age, is not well understood but could result from changes in immune system, microbiota, or diet, all of which change dramatically in early life.

Interestingly, the increase in resistance to *Cryptosporidium* infection correlates with time of weaning, when drastic shifts in the diversity and composition of the gut microbiota occur in both neonatal mice and human infants(10, 11). As enteric pathogens, *Cryptosporidium* primarily infect the apical end of small intestinal enterocytes, where they are enveloped by host membranes but remain extra-cytoplasmic (12, 13). Protrusion of the parasite-containing vacuole into the intestinal lumenal space places them near the mucosal layers and associated gut microbiota. In fact, several studies have shown that *C. parvum* infection alters the microbiota of mice(14, 15), and treatment with a probiotic enhanced *C. parvum* infection, presumably by altering the microbiota(16). Furthermore, loss of the microbiota in gnotobiotic and antibiotics-treated adult mice results in an increased susceptibility to *Cryptosporidium* infection(17), indicating that a diverse, mature microbiota provides a protective effect against *Cryptosporidium*. A recent study comparing different antibiotics revealed that cloxacilin treatment of mice induced changes in the microbiota and altered metabolites with increased susceptibility (18).

Since *Cryptosporidium* spends most of its life cycle inside a host cell, interactions between the parasite and the microbiota are likely mediated through metabolites in the intestinal lumenal space. Consistent with this idea, one study showed that high levels of fecal indole, a microbial metabolite, protected human volunteers from infection by *C. hominis* as monitored by oocysts shedding(19). While indole appears to inhibit the parasite, it is possible that other gut metabolites may promote *Cryptosporidium* growth. The genomes of *C. parvum*(20) and *C. hominis*(21) are highly streamlined, with the loss of many metabolic pathways and the expansion of transporters(22); hence, they must acquire many basic nutrients from their host or surrounding environs. It is possible that metabolites highly enriched in the neonatal gut, either derived from diet or the microbiota, are beneficial to the parasite and that the transition from milk to solid food, which is accompanied by changes in the microbiota, deprives *Cryptosporidium* of an essential nutrient.

In the present study, we undertook a systematic study of the changes in susceptibility of neonatal mice and the correlated change in the collective metabolites found in the lumen of the gut on the growth of *C. parvum*. Our findings reflect both enhancing and inhibitor activities of metabolites, indicating that gut metabolites influence susceptibility to infection during early development.

## Results

### Age-dependent susceptibility to *C. parvum* in a neonatal mouse model of cryptosporidiosis

To identify gut metabolites that may facilitate *Cryptosporidium* infection, we first determined the critical window of susceptibility to *C. parvum* in a neonatal mouse model of cryptosporidiosis. Four groups of ten pups each were reared simultaneously, and a subset of pups was infected each week with 5 × 10^4^ *C. parvum* oocysts (Fig. 1a, Supplementary Fig. 1). After five days of infection, the number of *C. parvum* genome equivalents in whole intestines was measured using quantitative PCR (qPCR) and normalized to the initial weight of the intestinal sample (Fig. 1b). Mice infected at one week of age had the highest number of *C. parvum* per gram of intestine, while parasite numbers dropped 10-fold in mice infected at two-weeks old (Fig. 1b). Mice inoculated at 3-weeks-old had the sharpest decline in *C. parvum* infection, with five orders of magnitude less *C. parvum* per gram of intestine than 1-week-old mice (Fig. 1b). Infection levels remained consistently lower for mice infected at 4, 5 and 6 weeks of age (Fig. 1b), indicating that mice are most susceptible to *C. parvum* infection within the first two weeks of life and experience a drastic reduction in parasite load when infected after this brief window of susceptibility.

**Fig 1.**
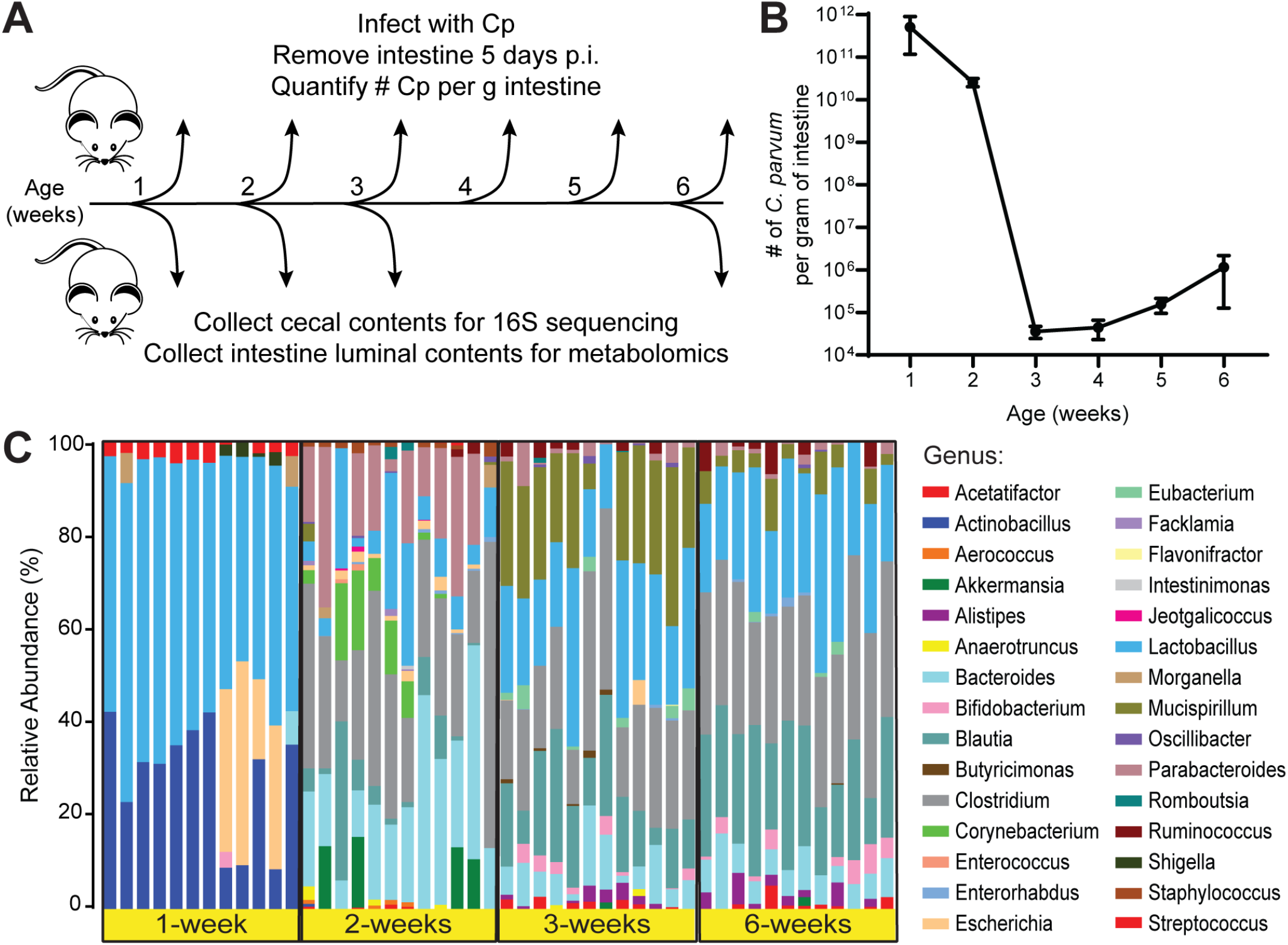
Differences between *C. parvum* infectivity and cecal microbiota during murine postnatal development. (A) Diagram of the experimental design. Separate cohorts of mice were challenged with 5 × 10^4^ *C. parvum* (Cp) oocysts at each week of life (N = 5-10 mice per week). Five days post infection (p.i.), intestines were removed, and the number of *C. parvum* organisms per gram of intestine was quantified by qPCR. In a separate experiment, cecal contents and small intestinal luminal contents were collected from uninfected mice at 1, 2, 3, and 6 weeks of age (N = 12 mice per week) for 16S ribosomal RNA sequencing and metabolomics, respectively. (B) Line graph depicting the average number of *C. parvum* organisms per gram intestine of mice infected at the indicated weeks of age (mean ± S.D., N = 10 mice each for weeks 1 and 2, N = 5 mice each for weeks 3-6). (C) Taxonomic differences in the cecal microbiota of mice at 1, 2, 3, or 6-weeks of age displayed as a stacked bar graph of the relative abundances of the bacterial genera detected by 16S ribosomal RNA sequencing.

To verify the course of age-dependent gut microbiome maturation in our model, we collected cecal contents from uninfected mice at timepoints when they are most susceptible (1 and 2 weeks of age) or relatively resistant (3 and 6 weeks of age) to infection (Fig. 1a, Supplementary Fig. 1) and performed 16s ribosomal RNA sequencing analysis. This analysis revealed drastic changes in the taxonomic composition of microbiota as the mice aged (Fig. 1c), similar to observations of previous studies in neonatal mice (10, 23–25). The microbial communities from 1-week-old mice were the least diverse of all four age groups (Supplementary Fig. 2a) and were dominated by facultative anaerobes from the Actinobacillus, Lactobacillus, and Escherichia genera (Fig. 1c, Supplementary Fig. 2b,c). By two weeks of age, the microbiota had transitioned to mostly strict anaerobes including Bacteroides, Parabacteroides, and Clostridium (Fig. 1c). In samples from 3-week and 6-week-old mice, Clostridium remained a significant fraction of the microbiota, while the relative abundances of Bacteroides and Parabacteroides decreased with a concurrent rise of the Blautia and Mucispirillum genera (Fig. 1c, Supplementary Fig. 2b,c). When all four age groups were analyzed together, a PCoA plot of weighted Unifrac distances shows distinct clusters for 1- and 2-week-old samples, while samples from 3- and 6-week-old mice overlap (Fig. 2a).

**Fig 2.**
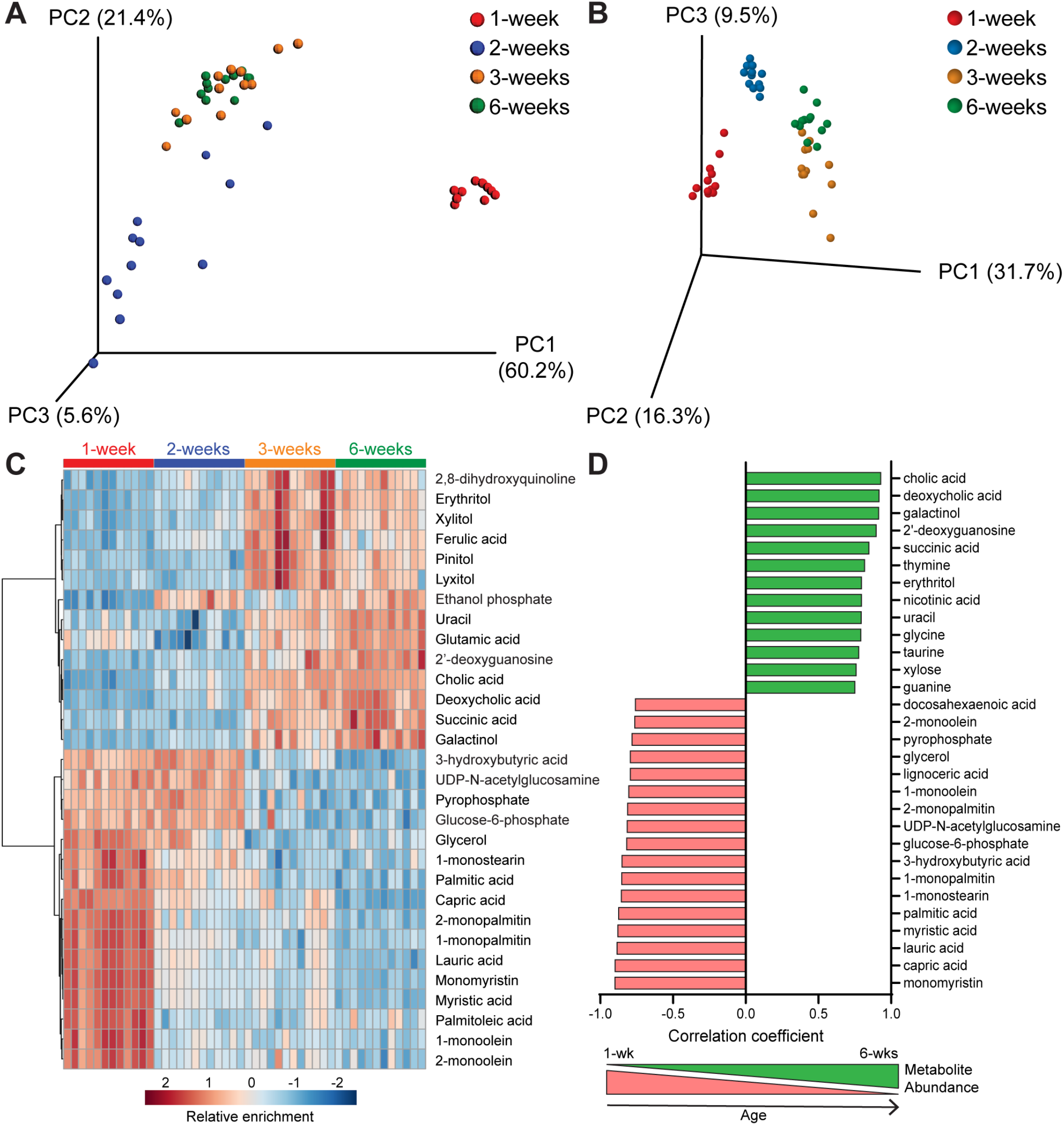
Differences in small intestinal metabolites during murine postnatal development. (A) PCoA plot of weighted Unifrac distances between cecal microbiota samples and (B) PCA plot of small intestinal metabolite differences from the same mice sampled at 1, 2, 3, or 6-weeks of age. (C) Hierarchical clustering of the top 30 metabolites that most significantly differed between groups as analyzed by one-way ANOVA, represented as a heat map with red indicating relative enrichment and blue indicating relative de-enrichment for the listed metabolites. (E) Bar graph showing the top 30 metabolites with relative abundances most significantly correlated with age by Pearson’s correlation. Red = negative correlation of metabolite abundance with age. Green = positive correlation of metabolite abundance with age.

### Changes in luminal metabolite composition over the first six weeks of life

To identify metabolites that could influence susceptibility to *C. parvum* infection, we collected small intestine luminal flush samples from the same mice as the microbiome analysis and quantified metabolites present using gas chromatography time-of-flight mass spectrometry (GC-TOF MS). A PCA plot of metabolite similarities between all samples revealed a similar pattern as the microbiome Unifrac analysis: metabolites from 1- and 2-week-old mice formed independent clusters, while those from 3- and 6-week-old mice were interspersed (Fig. 2b). Hierarchical clustering of the 30 metabolites with the lowest FDR-corrected p-values by one-way ANOVA revealed a strong enrichment of fatty acids and their glycerol esters (e.g., myristic acid and monomyristin; palmitic acid and monopalmitin) in 1-week samples only (Fig. 2c). In contrast, several metabolites, such as 3-hydroxybutric acid, UDP-N-acetylglucosamine and glucose-6-phosphate, were enriched in the first two weeks of life but decreased by three weeks. As expected given their overlapping PCA clusters (Fig. 2b), 3-week and 6-week samples were mostly enriched for the same metabolites (Fig. 2c) when compared to earlier timepoints. However, sugar alcohols such as erythritol, xylitol and lyxitol were generally more abundant at 3-weeks than at 6-weeks, while amino acids uracil and glutamic acid and bile acids (cholic and deoxycholic acid) were highest at 6-weeks (Fig. 2c).

A similar, but not identical, pattern emerged when Pearson’s correlation was used to find the top 30 metabolites whose abundances changed linearly over time (i.e., were either positively or negatively correlated with age) (Fig. 2d). The same fatty acids and their glycerol esters that were enriched in 1-week-old samples (Fig. 2c) were negatively correlated with age, with the addition of docosahexaenoic acid and lignoceric acid (Fig. 2d). Similarly, many of the metabolites enriched at the two later timepoints were positively correlated with age, with the cholic and deoxycholic bile acids having the strongest correlation (Fig. 2d).

### Screening for effects of neonatal metabolites on *C. parvum* growth in vitro

To determine if any of the metabolites negatively correlated with age (i.e., highest in 1-week-old samples) were sufficient to enhance *C. parvum* infection, we screened 43 metabolites for their effect on *C. parvum* growth in an HCT-8 human adenocarcinoma cell line (Table S1). All metabolites with a negative correlation with age (i.e. Pearson test) with an FDR-corrected *P* value < 0.05 were included in the screen except for those that proved insoluble or were not readily available for purchase. We also excluded metabolites associated with the microbiota of adult mice that were identified in a previous study comparing germ-free to recolonized mice (26). *C. parvum* growth was quantified using an image-based assay in which *C. parvum* oocysts were added with a single metabolite to HCT-8 cells plated in a 96-well format. After 24 hr of incubation, fixed cells were labeled with Pan-Cp, a polyclonal antibody that recognizes all stages of *C. parvum* (27), and stained with Hoechst 33342 to visualize host nuclei. The number of *C. parvum* and host nuclei in each well were quantified by an automated imaging platform and normalized to DMSO-treated control wells. Most metabolites were screened at 0.5 mM, but several metabolites required lower concentrations (either 0.1 mM or 0.02 mM) to avoid host toxicity issues (Fig. 3).

**Fig 3.**
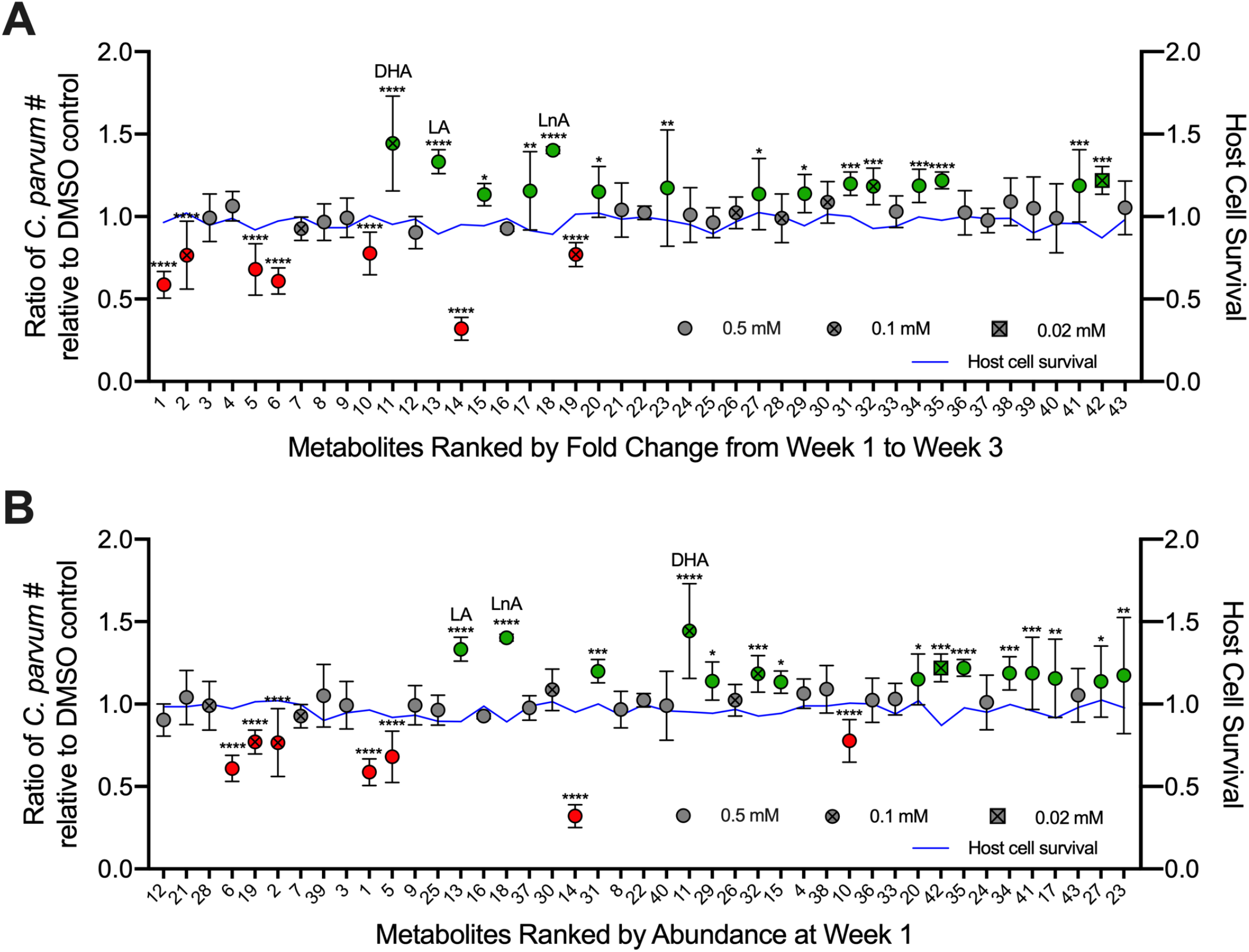
Effects of neonatal metabolites on *Cryptosporidium* growth. Average ratio of *C. parvum* parasites in treated samples relative to DMSO controls 24 hpi with metabolites in decreasing order of (A) fold decrease in abundance from mice aged 1 week to mice aged 3 weeks and (B) decreasing order of abundance in mice aged 1 week. Metabolites in green were found to significantly enhance growth, and metabolites in red were found to significantly inhibit growth. The three metabolites with the highest fold enhancement of growth are labeled: docosahexaenoic acid (DHA), linoleic acid (LA), and linolenic acid (LnA). * *P* ≤ 0.05, ** *P* ≤ 0.01, *** *P* ≤ 0.001, **** *P* ≤ 0.0001.

Out of the 43 metabolites screened, seven significantly inhibited *C. parvum* growth, while 15 significantly enhanced *C. parvum* infection compared to the DMSO control (Fig. 3). Interestingly, all of the inhibitory metabolites were medium- or long-chain saturated fatty acids and/or their glycerol esters: capric acid (C10:0); lauric acid (C12:0); myristic acid (C14:0) and monomyristin; palmitic acid (C16:0) and 1-monopalmitin; and 1-monostearin (C18:0). Not all saturated fatty acids were inhibitory: most had no effect and two, pentadecanoic acid (C15:0) and behenic acid (C22:0), modestly enhanced *C. parvum* growth. However, the three most potent enhancers (1.3 – 1.4X growth) were omega-3 or omega-6 polyunsaturated fatty acids: docosahexaenoic acid (DHA, C22:6), linolenic acid (LnA, C18:3) and linoleic acid (LA, C18:2). All the inhibitors and the most effective enhancers (DHA, LA, and LnA) fall within the top 20 metabolites when ranked based on their abundance fold change from week 1 to week 3 (Fig. 3a). When ranked by abundance at week 1, LA and LnA remain in the top 20 metabolites along with all inhibitors except for 1-monostearin (Fig. 3b). Thus, metabolites may have both protective and detrimental effects on susceptibility to *C. parvum* infection in the neonatal gut.

### Effects of omega-3 and omega-6 polyunsaturated fatty acids on *C. parvum* growth and invasion

Because the most potent enhancers in our screen were all omega-3 or omega-6 polyunsaturated fatty acids, we investigated whether other members of the omega-3 and omega-6 fatty acid families could positively affect *C. parvum* growth. Indeed, omega-3 eicosapentaenoic acid (EPA, C20:5) and omega-6 arachidonic acid (AA, C20:4) significantly enhanced *C. parvum* growth to the same extent as DHA, LA and LnA in a 24 hr growth assay (Fig. 4a), indicating that these two classes of fatty acids have a general positive effect on *C. parvum* infection. To investigate whether omega-3 and omega-6 fatty acids affect invasion efficiency of *C. parvum*, we infected HCT-8 cells with filtered sporozoites and treated with either DHA, LA, or LnA during a 2.5 hr invasion period. Cells were then extensively washed before fixing, staining, and imaging as described above. All three metabolites significantly increased the number of *C. parvum* present in the wells compared to the DMSO control, with LA and LnA having a slightly stronger effect than DHA (Fig. 4b). In contrast, parasite numbers did not significantly increase in HCT-8 cells that had been pretreated with DHA, LA, or LnA for 2 hr before infection with filtered sporozoites (Fig. 4b). This suggests that the fatty acids may be directly facilitating sporozoite adhesion or invasion to host cells, rather than acting through a host signaling pathway to “prime” the cells for invasion.

**Fig 4.**
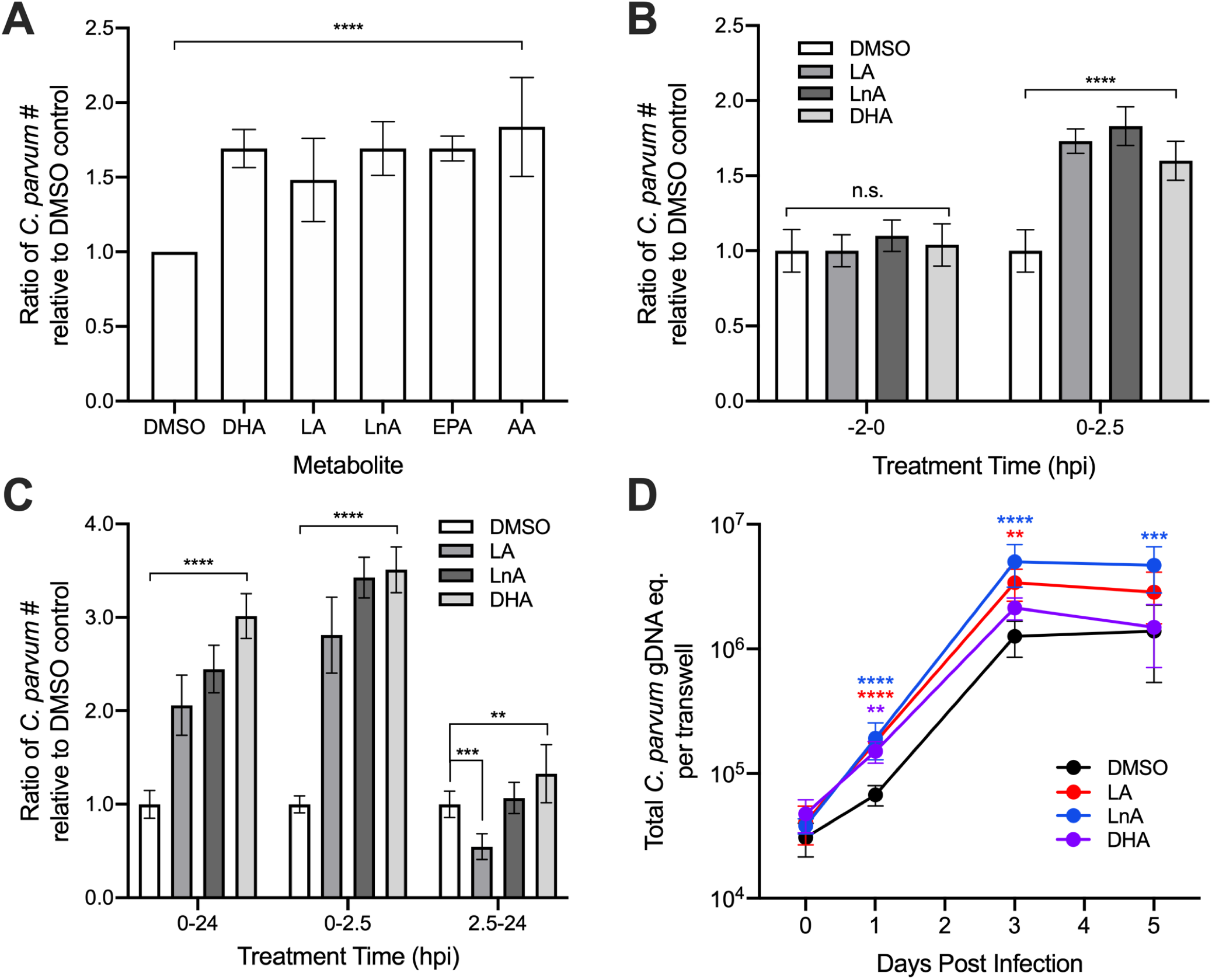
Enhancement of parasite growth and invasion by metabolites and related molecules. (A) Average ratio of *C. parvum* parasites 24 hpi in treated samples relative to DMSO controls for the metabolites docosahexaenoic acid (DHA), linoleic acid (LA), and linolenic acid (LnA) and the related compounds eicosapentaenoic acid (EPA) and arachidonic acid (AA). All tested compounds displayed significant enhancement of *C. parvum* growth compared to the DMSO control. (B) Samples were infected with filtered *C. parvum* sporozoites and washed after a 2.5 hr incubation period. Average ratio of attached *C. parvum* parasites relative to DMSO controls compared between samples that were pre-treated with metabolites for 2 hr and samples with metabolites added immediately after infection. (C) Average ratio of *C. parvum* parasites relative to DMSO control in samples infected with filtered sporozoites and treated with metabolites during invasion (0-2.5 hpi), after invasion (2.5-24 hpi), and for the duration of the experiment (0-24 hpi). (D) Effects of metabolite treatment on *C. parvum* growth in air-liquid interface (ALI) culture determined by average total *C. parvum* genomic DNA (gDNA) equivalent per transwell on days 0, 1, 3 and 5 post infection (mean ± S.D., N = 9 transwells per metabolite or N= 8 for DMSO). DHA and AA were tested at a final concentration of 0.1 mM and all other metabolites were tested at a final concentration of 0.5 mM. **** *P* ≤ 0.0001

To determine if the effects of metabolites on parasite growth may be time dependent, samples infected with filtered sporozoites were treated with LA, LnA, or DHA either during invasion (0-2.5 hours post-infection (hpi)), after invasion (2.5-24 hpi), or for the duration of the experiment (0-24 hpi). All samples were washed extensively 2.5 hpi to remove unattached sporozoites, and the culture media was replaced with or without metabolite solution depending on the respective treatment group. After 24 hpi, all samples were fixed, stained, and imaged as described above. For samples treated with LA, LnA, or DHA, parasite growth was significantly enhanced compared to the DMSO control when cells were treated from either 0-2.5 hpi or 0-24 hpi (Fig. 4c). However, when treatment began after invasion, treatment with LA significantly inhibited parasite growth, while treatment with LnA had no effect (Fig. 4c). Treatment with DHA from 2.5-24 hpi increased parasite growth relative to the control, but the magnitude of growth enhancement was far lower in samples treated after invasion than in samples where treatment began 0 hpi (Fig. 4c). These results indicate that the enhancement of parasite growth resulting from treatment with LA, LnA, and DHA is dependent on the presence of these metabolites during the first 2.5 hpi. This result implies that the increased parasite growth observed at later timepoints may directly result from the positive effects of metabolite treatment on sporozoite adhesion or invasion.

Because long-term culture and sexual reproduction of parasites is not supported in HCT-8 cell cultures, we tested whether metabolite treatment of parasites grown in air-liquid interface culture, a mouse ileal stem cell culture that allows complete development of the life cycle in vitro, would result in similar parasite growth enhancement(27). To determine this, transwells containing differentiated mouse intestinal epithelial cells (mIEC) were infected with filtered parasites and treated with LA, LnA, or DHA in both the top and bottom of transwells for 3 hr. All transwells were then washed to remove unattached sporozoites, and both the top and bottom of each transwell were treated with medium containing either DMSO or metabolite solution for the duration of the experiment. On days 0, 1, 3 and 5 post infection, DNA samples were collected from transwells, and *C. parvum* and mIEC genomic DNA quantities were determined using qPCR and standard curve analysis. Treatment with LA or DHA significantly enhanced parasite growth relative to the DMSO control at multiple timepoints, and treatment with LnA significantly enhanced the magnitude of parasite growth at all time points compared to the control (Fig. 4d). Metabolite treatments did not have adverse effects on epithelial culture and although they enhanced cell monolayer densities at some time points, this pattern did not correlate the enhanced growth of *C. parvum* (Fig. S1). Interestingly, treatment with LnA also increased the rate of parasite growth from day 0 to day 5 by 3-fold relative to the DMSO control, suggesting that the fatty acid may also be enhancing parasite replication after invasion. As a result, transwells treated continuously with LnA contained significantly greater quantities of C. parvum five days post infection (Fig. 4d).

## Discussion

Neonatal animals, including humans, are highly susceptible to *Cryptosporidium* infection, but quickly become resistant to the parasite as they age. In a neonatal mouse model of cryptosporidiosis, we found that susceptibility to the pathogen decreases sharply between two and three-weeks of life, concurrent with the cessation of breastfeeding and transition to solid food. This change in diet correlated with drastic shifts in the gut microbiota and lumenal metabolites, particularly the reduction of fatty acids typically found in breast milk. Exogenous addition of these fatty acids to in vitro cultures revealed that medium-to-long chain saturated fatty acids tend to inhibit *C. parvum* growth, while omega-3 and omega-6 polyunsaturated fatty acids enhance parasite invasion.

Previous studies in mice demonstrate that the gut microbiota changes dramatically during the first few weeks of life, especially following the dietary transition from breast milk to solid food(10, 24, 25). Specifically, these studies found that neonatal mice were first colonized by facultative anaerobes γ-Proteobacteria and Lactobacillales, which were progressively replaced by obligate anaerobes Clostridia and Bacteroidia during and after weaning(10, 24, 25). We observed similar developmental changes in the microbiota of our neonatal mice: in the first week of life, Lactobacillus and Actinobaccilus (a γ-proteobacteria) dominated the community and were replaced by two weeks of age with strict anaerobes including Clostridium and Bacteroides. Clostridium remained a significant fraction of the microbial community in mice post-weaning, while Bacteroides declined over time. Interestingly, a previous study that colonized germ free mice with cecal contents from neonatal (4-12 day) or adult mice (7 weeks) found that Clostridia (but not Bacteroides) protected mice colonized with adult microbiota against the enteric pathogens *Salmonella typhimurium* and *Citrobacter rodentium*(25). Although the protective mechanism is not fully understood, it was independent of innate and adaptive immune responses but was modulated by metabolites including succinate(25).

Concurrent with the microbial changes over time, the gut metabolome in our mice also transitioned as they aged: medium- and long-chain fatty acids were abundant in 1- and 2-week-old mice and were gradually replaced with sugar alcohols, amino acids, and bile salts in the 3- and 6-week-old mice. The abundance of fatty acids in pre-weaned mice reveals the significant contribution of diet to the overall gut metabolome, as fatty acids are important constituents of breast milk(28–31). In contrast, the metabolites enriched by week 6 begin to resemble those found in adult mice(26), and several are metabolic byproducts of intestinal bacteria, such as 2,8-hydroxyquinoline(32) and the secondary bile acid, deoxycholic acid(33). Hence, the shift in metabolite profiles after weaning is likely due to the absence of milk as a nutrient source as well as the production or induction of metabolites by a more mature microbiota.

Importantly, several of the fatty acids that were abundant in 1-week-old neonatal mice enhanced or inhibited the growth of the *C. parvum* in vitro. *C. parvum* is thought to lack a system for de novo fatty acid synthesis, instead relying on salvage from the host(22). It also lacks β-oxidation and cannot use fatty acids as an energy source(22). However, it contains three isoforms of the enzyme for acyl-Co-A addition (acyl co-A synthase (ACS))(34) needed for activating fatty acids salvaged from the host, a fatty acid synthase (FAS1) that functions as an elongase(35), and a long chain fatty acid elongase (LCE)(36). Of these enzymes, ACS isoforms prefer saturated substrates of C12-C18, and LCE prefers saturated substrates of C14-C16(36). The loading domain of CpFAS1 prefers palmitic acid (C16:0), but enzyme activity has been documented with substrates C12-C24 as well(37). Given these substrate preferences, it is somewhat surprising that medium chain fatty acids such as lauric acid (C12:0), myristic acid (C14:0) and palmitic acid (C16:0) inhibited *C. parvum* growth in vitro. One potential explanation for their inhibitory effects could be if these medium chain fatty acids inhibit the terminal reductase domain of FAS1, which normally prefers much longer chain substrates (i.e. > C24)(35). It is also possible that these fatty acids integrate into parasite membranes and upset the normal balance of lipids, hence compromising cellular functions. These medium-chain fatty acids have also been shown to inhibit bacterial growth in vitro(38, 39). Among these inhibitory compounds, capric acid, the strongest growth inhibitor in the 24 hr growth assay, has also been shown to inhibit *Candida albicans* growth and biofilm formation by altering gene expression(40), suggesting it has broad cross-phylum activity.

Several metabolites found early in the neonatal period enhanced *C. parvum* growth in vitro. For example, the long-chain, omega-3 or omega-6 polyunsaturated fatty acids, linoleic acid (LA, C18:2), linolenic acid (LnA, C18:3), and docosahexaenoic acid (DHA, C22:6) were all significant enhancers of parasite growth. Interestingly, this enhancement was also dependent on the timing of exposure: although pretreatment of host cells had no effect on subsequent infection, exposure during the first 2.5 hr of infection was critical to the enhancing effect. This timing suggests that these metabolites act to enhance invasion and/or formation of the parasitophorous vacuole that encases the parasite(12, 13). Since invasion and vacuole membrane formation require reorganization of host and parasite membranes in a rapid process of envelopment(41–43), the enhancing effects of these long chain unsaturated fatty acids may reflect the important properties they have on membrane composition, fluidity, and signaling(44, 45).

Although our studies were performed largely in vitro, they have important implications for infections in vivo. When metabolites were ranked by on overall abundance (based on the MS spectral counts), several of the enhancing metabolites, including LA, LnA, and DHA, were among the top half of the most abundant metabolites enriched in neonatal (1 and 2-week-old) pups, suggesting that they may contribute directly to increased susceptibility to *C. parvum* infection. Although our studies support a role for gut metabolites in modulating *C. parvum* infection, they are not the sole factors that determine increased susceptibility of neonatal mice to pathogens. Previous studies also indicate an important role for maturation of the immune system in susceptibility to infection during early life. In particular, CD103^+^ CD11c^+^ dendritic cells are found at low levels in neonatal mice and increase with maturation and during infection. Selective depletion of CD103+ dendritic cells in Batf3 knockout mice(46), or increase in their number by delivery of Flt-3 ligand(47), suggest that changes in these innate immune cells may underlie changes in susceptibility to *C. parvum* during maturation. Interestingly, administration of poly-IC to neonatal mice stimulated immune responses, including expanded DC cell functions, that required the presence of gut flora(48), indicating that the microbiota and immune function are tightly linked during early development.

The findings of our studies also have important implications for human cryptosporidiosis. The human microbiota undergoes similar predictable transitions from facultative aerobic bacteria such as Enterobacteriaceae at birth, to organisms that specialize on a milk-based diet such as Lactobacillus, then finally to a more mature, “adult-like” microbiota by 2 to 3-years of age(49–51). Interestingly, the microbiotas of children breast-feeding at 12-months-old are still dominated by Bifidobacterium and Lactobacillus, while the microbiotas of children that have stopped breast-feeding by this age are enriched in species prevalent in adults such as Clostridia(49). This suggests that the main driver of microbiota maturation is the cessation of breast-feeding and highlights the importance of breast milk in shaping the overall gut microbiota and metabolome. In our mice, polyunsaturated fatty acids in the gut lumen decreased significantly following weaning. Although fatty acid profiles in breast milk vary between species, all mammals produce essential fatty acids LA and LnA in their breast milk, as well as significant amounts of long-chain unsaturated fatty acids such as AA and DHA(31). Our finding that LA, LnA, AA and DHA all enhance sporozoite invasion suggests that human infants who are nursing may also be more susceptible to *Cryptosporidium* infection due to higher levels of these metabolites that are likely to be present in their guts, in comparisons to older, weaned children. These findings have important implications for the effects of diet and microbiota on the susceptibility of infants to cryptosporidiosis and possibly other enteric infections.

## Methods

### Neonatal mouse model of *C. parvum* infection

For infections of neonatal mice performed at the University of Arizona, *C. parvum* (Iowa strain) (52) oocysts were maintained by repeated passage in newborn *Cryptosporidium-free* Holstein bull calves (53), and purified from fecal material by sucrose density gradient centrifugation, as previously described (54).

To assess *C. parvum* infection levels with age in vivo, groups of 5 to 10 8-day-old specific pathogen-free ICR mice (Envigo) were used. All mice used in the present study were maintained in Biosafety Level 2 (BSL-2) biocontainment at the University of Arizona in accordance with the PHS *Guide for the Care and Use of Laboratory Animals* and IACUC approval.

Neonatal mice were randomly assigned to litters as detailed in **Figure S1**. At 1 week intervals after birth, mice were gavaged with 5 × 10^4^ *C. parvum* (Iowa strain) oocysts (N = 10 mice each for 1 and 2-weeks of age, N = 5 mice each for 3-6 weeks of age). At 5 days post-infection (92-94 hr post-infection), the entire intestine was extracted from each mouse, weighed, and then homogenized using ceramic beads in the Bead Ruptor4 (OMNI International, Kennesaw, GA). DNA was extracted using the QIAamp Fast DNA stool mini kit (Qiagen, Gaithersburg, MD) with the following modifications: after the addition of InhibitEx buffer, the samples were incubated at 95°C (5 min), followed by 5 freeze-thaw cycles using liquid nitrogen and a 37°C water bath. Total DNA in the samples was quantified by Nanodrop (Thermo Scientific, Waltham, MA).

Quantitative PCR (qPCR) for the *C. parvum* 18s rRNA (18S) was performed using the following primers: ChvF18S (5’-CAATAGCGTATATTAAAGTTGTTGCAGTT-3’ and ChvR18S (5’-CTGCTTTAAGCACTCTAATTTTCTCAAA-3’) (55)For qPCR, each 25 μL reaction contained a final concentration of 100 nM for both forward and reverse primers, (Invitrogen, Grand Island, NY) and 12.5 μL SYBR green Fast mix (Quantabio, Gaithersburg, MD). Genomic DNA (2 μL) was added, and the qPCR was performed in an ABI StepOne Plus Real-Time PCR System (Applied Biosystems, Grand Island, NY) with cycling conditions of 10 min incubation at 94°C followed by 45 cycles at 94°C for 10 sec, 54°C for 30 sec, and 72°C for 10 sec. Each sample was run in triplicate. A control with no template was run concurrently and was consistently negative. The number of *C. parvum* genomic equivalents was calculated for each sample based on a standard curve using DNA from known quantities of *C. parvum* oocysts and divided by the original weight of the intestinal sample to obtain the number of *C. parvum* organisms per gram intestine.

### Sample collection for 16S sequencing and metabolomics

Six pregnant ICR dams with litter sizes of 10 pups each were obtained from the same source (Envigo) as used for the neonatal infection experiment. Dams and the resulting pups were maintained in a specific pathogen-free barrier facility at Washington University School of Medicine with a strict 12-hr light cycle and ad libitum access to food and water. Mice were housed in complete autoclaved cage assemblies containing the same chow (Envigo NIH-31 Irradiated Modified Open Formula Mouse/Rat Diet 7913) and bedding (Envigo Teklad 7097 1/4” Corncob bedding) used in the neonatal infection experiment. To minimize experimental variation that could potentially arise from single cages or dams, 2 pups were randomly selected from each litter per timepoint (total of n=12 per timepoint) at 1-week, 2 weeks, 3 weeks, 4 weeks, and 6 weeks of age. Weaning was performed as usual at 3 weeks of age, with pups of the same sex housed only with littermates in fresh autoclaved cage assemblies. All procedures were approved by the Institutional Animal Care and Use Committee at Washington University School of Medicine. For collection of small intestinal luminal flushings for metabolomics, pups were euthanized and then the entire length of small intestine was dissected out intact and flushed with 500 uL of sterile phosphate-buffered saline (PBS) using a 1-mL syringe tipped with a blunt needle; small intestinal luminal contents from the flush were collected directly into a tared cryotube, weighed, and snap frozen in liquid nitrogen. For collection of cecal contents for 16s rRNA sequencing, the intact cecum was dissected and placed into a tared cryotube (pups aged 1, 2, or 3 weeks) or cecal contents were collected using a sterilized spatula and placed into a tared cryotube (pups aged 4 or 6 weeks); the material was weighed and then snap frozen in liquid nitrogen.

### 16S sequencing and analysis

DNA from cecal contents was isolated using the QIAamp DNA Stool Mini Kit (QIAGEN). The Washington University Genome Technology Access Center performed PCR amplification of all nine 16S variable regions with the Fluidigm Access Array System, indexing, pooling, and sequencing with an Illumina MiSeq Sequencer, as previously described(56). Sequencing data analysis either used the V1-V9 regions and the MVRSION pipeline(57) or the V4 region and QIIME pipeline version 1.9.0(58), as previously described(56). The OTU table resulting from QIIME analysis was used as input for linear discriminant analysis (LDA) effect size (LEfSe)(59) (http://huttenhower.sph.harvard.edu/lefse/) to identify statistically significant, differentially abundant taxa between the 1-week and 6-week old mice.

### Metabolite profiling and analysis

Untargeted metabolomics of the small intestinal luminal flushing samples by GC-TOF mass spectrometry was performed by the West Coast Metabolomics Center using the primary metabolism platform and a Leco Pegasus IV mass spectrometer. Of the 759 metabolites identified, 213 were annotated and used for further analysis. Data was normalized across samples by averaged Week1 values, before being log_2_ transformed and autoscaled. Data normalization and downstream univariate, multivariate, and clustering analyses were performed with Metaboanalyst 3.0 (https://www.metaboanalyst.ca)(60).

### HCT-8 cell culture and infection

For in vitro infection studies in human cell lines, *C. parvum* (AUCP-1 strain) oocysts were obtained from the Witola lab at the University of Illinois at Urbana-Champaign, where they were maintained by repeated passage in male Holstein calves and purified from fecal material as previously described. Animal procedures were approved by the Institutional Animal Studies Committee at the University of Illinois at Urbana-Champaign (61). Purified oocysts were stored at 4°C in PBS plus 50 mM Tris and 10 mM EDTA (pH 7.2) for up to six months before use.

Human ileocecal adenocarcinoma cells (HCT-8 cells, ATCC CCL-244) were maintained in RPMI 1640 medium (Gibco, ATCC modification) supplemented with 10% fetal bovine serum. Cells were confirmed to be mycoplasma free with the e-Myco plus *Mycoplasma* PCR detection kit (Boca Scientific).

### *C. parvum* growth assay for initial metabolite screen

Metabolites were chosen for the screen based on a negative Pearson coefficient and an FDR p-value ≤ 0.05. Metabolites that were insoluble or not readily available for purchase were excluded. We also excluded metabolites that had previously been shown to be present in the gut metabolome of germ-free mice and, thus, not likely produced or induced by the microbiota (26). In total, we tested 43 metabolites for their effects on *C. parvum* growth (Table S1).

All metabolites (Sigma-Aldrich) were reconstituted as 100 mM stock solutions in DMSO with the following exceptions: glucose-6-phosphate was dissolved in filtered PBS, phosphoethanolamine and glycerol-alpha-phosphate were dissolved in filtered dH_2_O, and cholesterol and arachidic acid were dissolved in filtered ethanol. HCT-8 cells were plated at 2 × 10^5^ cells per well in 96-well optically-clear-bottomed plates (Greiner Bio-One) and infected with 1.2 × 10^4^ to 5 × 10^4^ *C. parvum* oocysts (AUCP-1 strain) per well after 24 hr of cell growth. Metabolites were diluted in culture medium and immediately added to the wells following the addition of oocysts for a final metabolite concentration of 0.02 mM to 0.5 mM (depending on the metabolite) and 1% DMSO (three technical replicate wells per metabolite). Infected control wells containing only 1% DMSO media were included on each plate. At 24 hr after infection, wells were fixed in 4% formaldehyde for 10 min, washed twice with PBS, and then permeabilized and blocked for 20 min in blocking buffer composed of 0.1% Triton X-100 and 1% bovine serum albumin (BSA) in PBS. *C. parvum* were labeled with polyclonal rabbit anti-Cp antibody (27) diluted 1:2000 in blocking buffer, followed by goat anti-rabbit Alexa Fluor 488 (1:1000, Thermo Fisher Scientific). Host nuclei were stained with Hoechst 33342 (5 μg/ml, Thermo Fisher Scientific).

Plates were imaged with a 10X objective on a BioTek Cytation 3 cell imager (9 images per well in a 3 × 3 grid). Gen5 software version 5.0.2 was used to quantify the total number of parasites (puncta in the GFP channel) and host cells (nuclei in the DAPI channel) in images from each well. Relative parasite growth and host cell viability for each metabolite was calculated as a ratio of the mean number of *C. parvum* parasites or host cells, respectively, in the treated versus DMSO control groups averaged across three independent experiments with three technical replicates per experiment. Statistical analyses were performed in GraphPad Prism 8 using a two-way ANOVA followed by a Dunnett’s test for multiple comparisons, in which each metabolite was compared to the DMSO control.

### *C. parvum* invasion assay

HCT-8 cells were plated at 2 × 10^5^ cells per well in 96-well optically-clear-bottomed plates (Greiner Bio-one) and cultured for 24 hr as described above. To determine the effect of metabolite treatment on host cells before the addition of parasites, metabolite solutions diluted in culture media were added to half of the plate for a final concentration of 0.1 mM to 0.5 mM (depending on the metabolite) and 0.5% DMSO for 2 hr then washed 3x with PBS. Bleached *C. parvum* oocysts (AUCP-1 strain) were excysted for 1 hr at 37°C in a 0.75% sodium taurocholate solution and passed through a 1 μm filter to remove unexcysted oocysts. All wells were infected with approximately 2 × 10^5^ excysted sporozoites. Metabolite solutions diluted in culture media were then added to the second half of the plate for a final concentration of 0.1 mM to 0.5 mM and 0.5% DMSO. Control wells containing only 0.5% DMSO culture media were included for each half of the plate at each time point. After 2.5 hr of infection, wells were fixed and stained with polyclonal rabbit anti-Cp antibody (1:5000), goat anti-rabbit Alexa Fluor 488 (1:1000, Thermo Fisher Scientific), and Hoescht 33342 (5 μg/ml, Thermo Fisher Scientific) as detailed above.

Parasites and host cells were imaged and quantified using the same protocol as the *C. parvum* growth assay. Relative parasite growth and host cell viability for each metabolite was calculated as a ratio of the mean number of *C. parvum* parasites or host cells, respectively, in the treated versus DMSO control groups averaged across three independent experiments with three technical replicates per experiment. Statistical analyses were performed in GraphPad Prism 8 using a two-way ANOVA followed by a Dunnett’s test for multiple comparisons, in which each metabolite was compared to the DMSO control within each treatment group.

### *C. parvum* invasion wash-out assay

HCT-8 cells were plated 2 × 10^5^ cells per well in a clear-bottomed 96 well plate. After 24 hr of cell growth, cells were infected with 1 × 10^5^ filtered, excysted sporozoites per well. Immediately after infection, metabolite solutions (0.5 mM except for DHA that was 0.1 mM) or DMSO control were added to wells. After 2.5 hr of incubation, all wells were washed 2x with PBS, and metabolite solutions or DMSO control were added to wells as appropriate for each group. At 24 hpi, all wells were fixed and stained as described for the *C. parvum* invasion assay above.

Parasites and host cells were imaged and quantified as detailed in the *C. parvum* growth assay. Relative parasite growth and host cell viability for each metabolite were calculated as a ratio of the mean number of *C. parvum* parasites or host cells, respectively, in the treated versus DMSO control groups averaged across three independent experiments with three technical replicates per experiment. Statistical analyses were performed in GraphPad Prism 8 using a two-way ANOVA followed by a Dunnett’s test for multiple comparisons, in which each metabolite was compared to the DMSO control within each treatment group.

### Quantification of *C. parvum* in metabolite-treated air-liquid interface transwells

Mouse intestinal epithelial cells (mIEC) monolayers were cultured on transwells with an air-liquid interface (ALI) as previously described(27, 62). Briefly, irradiated 3T3 mouse fibroblast cells (CRL-1658 ATCC) were plated on transwells (polyester membrane, 0.4 μm pore; Corning Costar) coated with 10% Matrigel (Corning) and cultured at 37 °C for approximately 24 hr in Dulbecco’s Modified Eagle’s Medium (DMEM high glucose; Sigma D6429) with 10% fetal bovine serum (Sigma) and 1X penicillin/streptomycin (Sigma). Primary mouse ileal stem cells were harvested from three-day old spheroid cultures in Matrigel, dissociated with trypsin as previously described(63), and plated onto irradiated i3T3 monolayers at 5×10^4^ mIECs per transwell. mIEC monolayers were cultured with 50% L-WRN conditioned medium(64) and 10 μM Y-27632 ROCK inhibitor (Torcis Bioscience) in both the top and bottom compartments of the transwell for 7 days, after which medium was removed from the top compartment to create the air-liquid interface. Three days after removing the top medium, each transwell was infected with 2 × 10^5^ filtered, excysted sporozoites, and DMSO control or metabolite solutions (0.5 mM except for DHA that was 0.1 mM) were added to both the top (50 μl) and bottom (400 μl) compartments of the transwell. After 3 hr of incubation, top medium was removed and each transwell was washed with PBS. Each transwell was then treated continuously with either DMSO control or metabolite solution in both the top and bottom chamber for the duration of the experiment.

DNA from transwells was collected and extracted using the QIAmp DNA Mini kit (QIAGEN). qPCR was performed using the QuantStudio 3 System with cycling conditions of a 10 min incubation at 95 °C, then 40 cycles at 95 °C for 15s and 60 °C for 1 min, followed by a continuous melt curve analysis to identify samples with evidence of non-specific amplification. Each reaction contained 2 μL purified transwell DNA (diluted 1:5) as a template, 10 μL SYBR Green QuickStart Taq ReadyMix (Sigma), and 1.6 μL of 5 μM primer solution targeting *C. parvum* GAPDH (forward: 5’-CGGATGGCCATACCTGTGAG-3’ and reverse: 5’-GAAGATGCGCTGGGAACAAC-3’)(27) or mouse GAPDH (forward: 5’-GCCATGAGTGGACCCTTCTT-3’ and reverse: 5’-GAAAACACGGGGGCAATGAG-3’)(27). Each transwell sample was run with technical duplicates, and negative (water) controls were included in each plate.

*C. parvum* and mIEC genomic DNA (gDNA) quantities per transwell were determined via the QuantStudio Design & Analysis New (DA2) software using standard curves for *C. parvum* and mouse gDNA, respectively. Total *C. parvum* or mIEC gDNA per transwell was calculated as an average of gDNA quantities per transwell across three independent experiments with two to three technical replicates per experiment. Statistical analyses were performed in GraphPad Prism 8 using a two-way ANOVA followed by a Dunnett’s test for multiple comparisons, in which each metabolite was compared to the DMSO control within each timepoint.

## Data Availability

16S ribosomal RNA sequencing reads are available in the ArrayExpress database (http://www.ebi.ac.uk/arrayexpress) under accession number E-MTAB-9100. All remaining data discussed in this report are found in the main figures or the supplementary materials.

## ACKNOWLEDGEMENTS

We are grateful to William Witola, University of Illinois, for providing the *C. parvum* oocysts used here, Megan Baldridge for helpful advice, and Soumya Ravindran for cell culture support. Supported in part by grants from the NIH (AI 145496 to L.D.S.). K.L.V. was supported by NIH grant DK109081. We thank the Genome Technology Access Center (GTAC) in the Department of Genetics at Washington University School of Medicine for help with 16s rRNA sequencing and analysis. The GTAC is partially supported by NCI Cancer Center Support Grant #P30 CA91842 to the Siteman Cancer Center and by ICTS/CTSA Grant# UL1TR002345 from the National Center for Research Resources (NCRR), a component of the National Institutes of Health (NIH), and NIH Roadmap for Medical Research. This publication is solely the responsibility of the authors and does not necessarily represent the official view of NCRR or NIH.

## Supplemental Material

Fig S1. Experimental timeline for testing murine susceptibility to *C. parvum* and for collecting samples for meta

Fig S2. Cecal microbiota taxonomic differences during murine postnatal development.

Fig S3. Average host cell viability of Air-liquid Interface (ALI) culture

## REFERENCES

1. Kotloff KL, Nataro JP, Blackwelder WC, Nasrin D, Farag TH, Panchalingam S, Wu Y, Sow SO, Sur D, Breiman RF, Faruque AS, Zaidi AK, Saha D, Alonso PL, Tamboura B, Sanogo D, Onwuchekwa U, Manna B, Ramamurth T, Kanungo S, Ochieng JB, Omore R, Oundo JO, Hossain A, Das SK, Ahmed S, Qureshi S, Quadri F, Adegbola RA, Antonio M, Hossain MJ, Akinsola A, Mandomando I, Nhampossa T, Acácio S, Biswas K, O’Reilly CE, Mintz ED, Berkeley LY, Muhsen K, Sommerfelt H, Robins-Browne RM, Levine MM. 2013. Burden and aetiology of diarrhoeal disease in infants and young children in developing countries (the Global Enteric Multicenter Study, GEMS): a prospective, case-control study. Lancet 382:209–22.

2. Kotloff KL. 2017. The Burden and Etiology of Diarrheal Illness in Developing Countries. Pediatr Clin North Am 64:799–814.

3. Fayer R. 2004. Cryptosporidium: a water-borne zoonotic parasite. Vet Parasitol 126:37–56.

4. Peng MM, Xiao L, Freeman AR, Arrowood MJ, Escalante AA, Weltman AC, Ong CSL, Mac Kenzie WR, Lal AA, Beard CB. 1997. Genetic polymorphism among *Cryptosporidium parvum* isolates: evidence of two distinct human transmission. Emerging Infectious Diseases 3:567–573.

5. Feng Y, Ryan UM, Xiao L. 2018. Genetic diversity and population structure of *Cryptosporidium*. Trends Parasitol 34:997–1011.

6. Feng Y, Tiao N, Li N, Hlavsa M, Xiao L. 2014. Multilocus sequence typing of an emerging Cryptosporidium hominis subtype in the United States. J Clin Microbiol 52:524–30.

7. Checkley W, White AC, Jr., Jaganath D, Arrowood MJ, Chalmers RM, Chen XM, Fayer R, Griffiths JK, Guerrant RL, Hedstrom L, Huston CD, Kotloff KL, Kang G, Mead JR, Miller M, Petri WA, Jr., Priest JW, Roos DS, Striepen B, Thompson RC, Ward HD, Van Voorhis WA, Xiao L, Zhu G, Houpt ER. 2015. A review of the global burden, novel diagnostics, therapeutics, and vaccine targets for cryptosporidium. Lancet Infect Dis 15:85–94.

8. Sherwood D, Angus KW, Snodgrass DR, Tzipori S. 1982. Experimental cryptosporidiosis in laboratory mice. Infect Immun 38:471–5.

9. Zambriski JA, Nydam DV, Bowman DD, Bellosa ML, Burton AJ, Linden TC, Liotta JL, Ollivett TL, Tondello-Martins L, Mohammed HO. 2013. Description of fecal shedding of Cryptosporidium parvum oocysts in experimentally challenged dairy calves. Parasitol Res 112:1247–54.

10. Al Nabhani Z, Dulauroy S, Marques R, Cousu C, Al Bounny S, Dejardin F, Sparwasser T, Berard M, Cerf-Bensussan N, Eberl G. 2019. A Weaning Reaction to Microbiota Is Required for Resistance to Immunopathologies in the Adult. Immunity 50:1276–1288 e5.

11. Subramanian S, Huq S, Yatsunenko T, Haque R, Mahfuz M, Alam MA, Benezra A, DeStefano J, Meier MF, Muegge BD, Barratt MJ, VanArendonk LG, Zhang Q, Province MA, Petri WA, Jr., Ahmed T, Gordon JI. 2014. Persistent gut microbiota immaturity in malnourished Bangladeshi children. Nature 510:417–21.

12. Current WL, Reese NC. 1986. A comparison of endogenous development of three isolates of Cryptosporidium in suckling mice. J Protozool 33:98–108.

13. Umemiya R, Fukuda M, Fujisaki K, Matsui T. 2005. Electron microscopic observation of the invasion process of Cryptosporidium parvum in severe combined immunodeficiency mice. J Parasitol 91:1034–9.

14. Ras R, Huynh K, Desoky E, Badawy A, Widmer G. 2015. Perturbation of the intestinal microbiota of mice infected with Cryptosporidium parvum. Int J Parasitol 45:567–73.

15. Mammeri M, Chevillot A, Thomas M, Julien C, Auclair E, Pollet T, Polack B, Vallee I, Adjou KT. 2019. Cryptosporidium parvum-Infected Neonatal Mice Show Gut Microbiota Remodelling Using High-Throughput Sequencing Analysis: Preliminary Results. Acta Parasitol 64:268–275.

16. Oliveira BCM, Widmer G. 2018. Probiotic Product Enhances Susceptibility of Mice to Cryptosporidiosis. Appl Environ Microbiol 84.

17. Harp JA, Wannemuehler MW, Woodmansee DB, Moon HW. 1988. Susceptibility of germfree or antibiotic-treated adult mice to Cryptosporidium parvum. Infect Immun 56:2006–10.

18. Charania R, Wade BE, McNair NN, Mead JR. 2020. Changes in the Microbiome of Cryptosporidium-Infected Mice Correlate to Differences in Susceptibility and Infection Levels. Microorganisms 8.

19. Chappell CL, Okhuysen PC, Langer-Curry R, Widmer G, Akiyoshi DE, Tanriverdi S, Tzipori S. 2006. Cryptosporidium hominis: experimental challenge of healthy adults. Am J Trop Med Hyg 75:851–7.

20. Abrahamsen MS, Templeton TJ, Enomoto S, Abrahante JE, Zhu G, Lancto CA, Deng M, Liu C, Widmer G, Tzipori S, Buck GA, Xu P, Bankier AT, Dear PH, Konfortov BA, Spriggs HF, Lakshminarayan I, Anantharaman V, Aravind L, Kapur V. 2004. Complete genome sequence of the apicomplexan, *Cryptosporidium parvum*. Sciencexpress 304:441–445.

21. Xu P, Widmer G, Wang Y, Ozaki LS, Alves JM, Serrano MG, Puiu D, Manque P, Akiyoshi D, Mackey AJ, Pearson WR, Dear PH, Bankier AT, Peterson DL, Abrahamsen MS, Kapur V, Tzipori S, Buck GA. 2004. The genome of Cryptosporidium hominis. Nature 431:1107–12.

22. Rider SD, Jr., Zhu G. 2010. Cryptosporidium: genomic and biochemical features. Exp Parasitol 124:2–9.

23. Singer JR, Blosser EG, Zindl CL, Silberger DJ, Conlan S, Laufer VA, DiToro D, Deming C, Kumar R, Morrow CD, Segre JA, Gray MJ, Randolph DA, Weaver CT. 2019. Preventing dysbiosis of the neonatal mouse intestinal microbiome protects against late-onset sepsis. Nat Med 25:1772–1782.

24. Knoop KA, Gustafsson JK, McDonald KG, Kulkarni DH, Coughlin PE, McCrate S, Kim D, Hsieh CS, Hogan SP, Elson CO, Tarr PI, Newberry RD. 2017. Microbial antigen encounter during a preweaning interval is critical for tolerance to gut bacteria. Sci Immunol 2.

25. Kim YG, Sakamoto K, Seo SU, Pickard JM, Gillilland MG, 3rd, Pudlo NA, Hoostal M, Li X, Wang TD, Feehley T, Stefka AT, Schmidt TM, Martens EC, Fukuda S, Inohara N, Nagler CR, Nunez G. 2017. Neonatal acquisition of Clostridia species protects against colonization by bacterial pathogens. Science 356:315–319.

26. Matsumoto M, Kibe R, Ooga T, Aiba Y, Kurihara S, Sawaki E, Koga Y, Benno Y. 2012. Impact of intestinal microbiota on intestinal luminal metabolome. Sci Rep 2:233.

27. Wilke G, Funkhouser-Jones LJ, Wang Y, Ravindran S, Wang Q, Beatty WL, Baldridge MT, VanDussen KL, Shen B, Kuhlenschmidt MS, Kuhlenschmidt TB, Witola WH, Stappenbeck TS, Sibley LD. 2019. A stem-cell-derived platform enables complete *Cryptosporidium* development in vitro and genetic tractability. Cell Host Microbe 26:123–134 e8.

28. Qian L, Zhao A, Zhang Y, Chen T, Zeisel SH, Jia W, Cai W. 2016. Metabolomic Approaches to Explore Chemical Diversity of Human Breast-Milk, Formula Milk and Bovine Milk. Int J Mol Sci 17.

29. Smilowitz JT, O’Sullivan A, Barile D, German JB, Lonnerdal B, Slupsky CM. 2013. The human milk metabolome reveals diverse oligosaccharide profiles. J Nutr 143:1709–18.

30. Oosting A, Verkade HJ, Kegler D, van de Heijning BJ, van der Beek EM. 2015. Rapid and selective manipulation of milk fatty acid composition in mice through the maternal diet during lactation. J Nutr Sci 4:e19.

31. Neville MC, Picciano MF. 1997. Regulation of milk lipid secretion and composition. Annu Rev Nutr 17:159–83.

32. Hubbard TD, Liu Q, Murray IA, Dong F, Miller C, 3rd, Smith PB, Gowda K, Lin JM, Amin S, Patterson AD, Perdew GH. 2019. Microbiota Metabolism Promotes Synthesis of the Human Ah Receptor Agonist 2,8-Dihydroxyquinoline. J Proteome Res 18:1715–1724.

33. Ridlon JM, Kang DJ, Hylemon PB. 2006. Bile salt biotransformations by human intestinal bacteria. J Lipid Res 47:241–59.

34. Guo F, Zhang H, Payne HR, Zhu G. 2016. Differential Gene Expression and Protein Localization of Cryptosporidium parvum Fatty Acyl-CoA Synthetase Isoforms. J Eukaryot Microbiol 63:233–46.

35. Zhu G, Shi X, Cai X. 2010. The reductase domain in a Type I fatty acid synthase from the apicomplexan Cryptosporidium parvum: restricted substrate preference towards very long chain fatty acyl thioesters. BMC Biochem 11:46.

36. Fritzler JM, Millership JJ, Zhu G. 2007. Cryptosporidium parvum long-chain fatty acid elongase. Eukaryot Cell 6:2018–28.

37. Zhu G, Li Y, Cai X, Millership JJ, Marchewka MJ, Keithly JS. 2004. Expression and functional characterization of a giant Type I fatty acid synthase (CpFAS1) gene from Cryptosporidium parvum. Mol Biochem Parasitol 134:127–35.

38. Huang CB, Alimova Y, Myers TM, Ebersole JL. 2011. Short- and medium-chain fatty acids exhibit antimicrobial activity for oral microorganisms. Arch Oral Biol 56:650–4.

39. Marounek M, Skrivanova E, Rada V. 2003. Susceptibility of Escherichia coli to C2-C18 fatty acids. Folia Microbiol (Praha) 48:731–5.

40. Jadhav A, Mortale S, Halbandge S, Jangid P, Patil R, Gade W, Kharat K, Karuppayil SM. 2017. The Dietary Food Components Capric Acid and Caprylic Acid Inhibit Virulence Factors in Candida albicans Through Multitargeting. J Med Food 20:1083–1090.

41. Chen XM, O’Hara SP, Huang BQ, Nelson JB, Lin JJ, Zhu G, Ward HD, LaRusso NF. 2004. Apical organelle discharge by Cryptosporidium parvum is temperature, cytoskeleton, and intracellular calcium dependent and required for host cell invasion. Infect Immun 72:6806–16.

42. Elliott DA, Clark DP. 2000. Cryptosporidium parvum induces host cell actin accumulation at the host-parasite interface. Infect Immun 68:2315–22.

43. Wetzel DM, Schmidt J, Kuhlenschmidt M, Dubey JP, Sibley LD. 2005. Gliding motility leads to active cellular invasion by *Cryptosporidium parvum* sporozoites. Infect Immun 73:5379–5387.

44. Stubbs CD, Smith AD. 1984. The modification of mammalian membrane polyunsaturated fatty acid composition in relation to membrane fluidity and function. Biochim Biophys Acta 779:89–137.

45. Schmitz G, Ecker J. 2008. The opposing effects of n-3 and n-6 fatty acids. Prog Lipid Res 47:147–55.

46. Potiron L, Lacroix-Lamande S, Marquis M, Levern Y, Fort G, Franceschini I, Laurent F. 2019. Batf3-Dependent Intestinal Dendritic Cells Play a Critical Role in the Control of Cryptosporidium parvum Infection. J Infect Dis 219:925–935.

47. Lantier L, Lacroix-Lamande S, Potiron L, Metton C, Drouet F, Guesdon W, Gnahoui-David A, Le Vern Y, Deriaud E, Fenis A, Rabot S, Descamps A, Werts C, Laurent F. 2013. Intestinal CD103+ dendritic cells are key players in the innate immune control of Cryptosporidium parvum infection in neonatal mice. PLoS Pathog 9:e1003801.

48. Lantier L, Drouet F, Guesdon W, Mancassola R, Metton C, Lo-Man R, Werts C, Laurent F, Lacroix-Lamande S. 2014. Poly(I:C)-induced protection of neonatal mice against intestinal Cryptosporidium parvum infection requires an additional TLR5 signal provided by the gut flora. J Infect Dis 209:457–67.

49. Backhed F, Roswall J, Peng Y, Feng Q, Jia H, Kovatcheva-Datchary P, Li Y, Xia Y, Xie H, Zhong H, Khan MT, Zhang J, Li J, Xiao L, Al-Aama J, Zhang D, Lee YS, Kotowska D, Colding C, Tremaroli V, Yin Y, Bergman S, Xu X, Madsen L, Kristiansen K, Dahlgren J, Wang J. 2015. Dynamics and Stabilization of the Human Gut Microbiome during the First Year of Life. Cell Host Microbe 17:852.

50. Bokulich NA, Chung J, Battaglia T, Henderson N, Jay M, Li H, A DL, Wu F, Perez-Perez GI, Chen Y, Schweizer W, Zheng X, Contreras M, Dominguez-Bello MG, Blaser MJ. 2016. Antibiotics, birth mode, and diet shape microbiome maturation during early life. Sci Transl Med 8:343ra82.

51. Yatsunenko T, Rey FE, Manary MJ, Trehan I, Dominguez-Bello MG, Contreras M, Magris M, Hidalgo G, Baldassano RN, Anokhin AP, Heath AC, Warner B, Reeder J, Kuczynski J, Caporaso JG, Lozupone CA, Lauber C, Clemente JC, Knights D, Knight R, Gordon JI. 2012. Human gut microbiome viewed across age and geography. Nature 486:222–7.

52. Heine J, Pohlenz JF, Moon HW, Woode GN. 1984. Enteric lesions and diarrhea in gnotobiotic calves monoinfected with Cryptosporidium species. J Infect Dis 150:768–75.

53. Riggs MW, McGuire TC, Mason PH, Perryman LE. 1989. Neutralization-sensitive epitopes are exposed on the surface of infectious Cryptosporidium parvum sporozoites. J Immunol 143:1340–5.

54. Arrowood MJ, Sterling CR. 1987. Isolation of Cryptosporidium oocysts and sporozoites using discontinuous sucrose and isopycnic Percoll gradients. J Parasitol 73:314–9.

55. Burnet JB, Ogorzaly L, Tissier A, Penny C, Cauchie HM. 2013. Novel quantitative TaqMan real-time PCR assays for detection of Cryptosporidium at the genus level and genotyping of major human and cattle-infecting species. J Appl Microbiol 114:1211–22.

56. Liu TC, Kern JT, VanDussen KL, Xiong S, Kaiko GE, Wilen CB, Rajala MW, Caruso R, Holtzman MJ, Gao F, McGovern DP, Nunez G, Head RD, Stappenbeck TS. 2018. Interaction between smoking and ATG16L1T300A triggers Paneth cell defects in Crohn’s disease. J Clin Invest 128:5110–5122.

57. Schriefer AE, Cliften PF, Hibberd MC, Sawyer C, Brown-Kennerly V, Burcea L, Klotz E, Crosby SD, Gordon JI, Head RD. 2018. A multi-amplicon 16S rRNA sequencing and analysis method for improved taxonomic profiling of bacterial communities. J Microbiol Methods 154:6–13.

58. Caporaso JG, Kuczynski J, Stombaugh J, Bittinger K, Bushman FD, Costello EK, Fierer N, Pena AG, Goodrich JK, Gordon JI, Huttley GA, Kelley ST, Knights D, Koenig JE, Ley RE, Lozupone CA, McDonald D, Muegge BD, Pirrung M, Reeder J, Sevinsky JR, Turnbaugh PJ, Walters WA, Widmann J, Yatsunenko T, Zaneveld J, Knight R. 2010. QIIME allows analysis of high-throughput community sequencing data. Nat Methods 7:335–6.

59. Segata N, Izard J, Waldron L, Gevers D, Miropolsky L, Garrett WS, Huttenhower C. 2011. Metagenomic biomarker discovery and explanation. Genome Biol 12:R60.

60. Xia J, Sinelnikov IV, Han B, Wishart DS. 2015. MetaboAnalyst 3.0--making metabolomics more meaningful. Nucleic Acids Res 43:W251–7.

61. Zhang X, Kim CY, Worthen T, Witola WH. 2018. Morpholino-mediated in vivo silencing of *Cryptosporidium parvum* lactate dehydrogenase decreases oocyst shedding and infectivity. Int J Parasitol 48:649–656.

62. Wilke G, Wang Y, Ravindran S, Stappenbeck T, Witola WH, Sibley LD. 2020. In vitro culture of *Cryptosporidium parvum* using stem cell-derived intestinal epithelial monolayers, p 351–372. *In* Mead J, Arrowood M (ed), Cryptosporidium Methods in Molecular Biology, vol 2052. Humana, New York, NY.

63. Moon C, VanDussen KL, Miyoshi H, Stappenbeck TS. 2014. Development of a primary mouse intestinal epithelial cell monolayer culture system to evaluate factors that modulate IgA transcytosis. Mucosal Immunol 7:818–28.

64. Miyoshi H, Ajima R, Luo C, Yamaguchi TP, Stappenbeck TS. 2012. Wnt5a potentiates TGF-beta signaling to promote colonic crypt regeneration after tissue injury.. Science 338:108–13.

